# Zombie Gene Flow: Asexual Hybrids Mediate Extensive Genomic Introgression from Extinct Species Into Their Sexual Parent

**DOI:** 10.1101/2025.03.28.645731

**Authors:** Jasna Vukić, Anatolie Marta, Filip Matura, Daniel Kulik, Jan Eisner, Radek Šanda, Tomáš Tichopád, Lukáš Choleva, Anabel Perdices, Jan Roslein, Jan Kočí, Spase Shumka, Denik Ulqini, Jan Kotusz, Dmitrij Dedukh, Karel Janko

**Author notes:** corresponding author: Karel Janko. co-corresponding author: Dmitrij Dedukh.

## Abstract

Interspecific gene flow may profoundly impact genome integrity and adaptive evolution in hybridizing species, leading to novelties such as transgressive traits, supergenes, or, sometimes, the emergence of asexually reproducing lineages. Conventionally, introgression is thought to proceed between reproductively interacting species, mediated by recombining interspecific hybrids, while asexual lineages are considered evolutionary deadlock for genomes trapped in them. Our study on *Cobitis* loaches in the western Balkan watersheds demonstrates an alternative mechanism where a mix of asexuality and polyploidy facilitates significant introgression from a long-extinct species. Through extensive sampling and cytogenetic and phylogenomic analyses, we identified a sexual species, *C. ohridana* (OO) coexisting with its asexual hybrid form (OX) originating from hybridization with an extinct species (XX). The diploid OX hybrids pass both parental subgenomes mostly clonally with occasional gene conversions, while triploid OOX hybrids reproduce through meiotic hybridogenesis, producing O-like gametes with localized gene conversions by X alleles. Their mating with the locally dominant sexual species consequently leads to over 4% admixture in both nuclear and mitochondrial genomes. Our findings challenge the view of hybrid asexual lineages as evolutionary dead ends, revealing their significant role as reservoirs of genetic diversity and agents of interspecific gene exchange, even after the extinction of one parental taxon.

## INTRODUCTION

Hybridization, the interspecific gene flow, is a potent evolutionary force, fostering diversifications and radiations across various animal and plant groups (Arnold 1996; Meier et al. 2023), and influencing genetic architectures of recipient. The transfer of beneficial alleles between species may enhance their adaptive potential (Brauer et al. 2023), and the introgression of diverged structural variants can constrain recombination, potentially leading to the formation of supergenes (Schwander et al. 2014). A crucial prerequisite for introgressive hybridization is the existence of fertile hybrids capable of backcrossing through recombinant gametes. However, the reproductive abilities of hybrids critically depend on the compatibility of parental genomes, which diminishes as species divergence progresses.

The commonly invoked speciation continuum concept (e.g. Seehausen et al. 2014) predicts quantitative reduction in fertility and viability of hybrids, effectively ceasing gene flow as divergence between parental species increases (e.g. Bolnick and Near 2005)). Yet, recent research demonstrated that increasing genetic divergence between species also implies a qualitative shift in hybrid’s reproductive strategies towards the production of unreduced gametes (Janko et al. 2018; Barley et al. 2022), thereby also paving the road towards formation of polyploid forms (Choleva and Janko 2013). Accumulation of incompatibilities, presumably in gene pathways regulating the cell cycle and meiosis (Moritz et al. 1989), thus may temporarily open a unique window along the so-called ***‘extended speciation continuum***’ (Stöck et al. 2021), where hybridization induces asexuality, which in turn acts as a speciation barrier preventing genetic recombination and backcrossing, while maintaining the fertility of hybrids. Asexual reproduction seen among natural hybrids takes various forms, such as complete alteration of meiosis through slippage of the first meiotic division (Dedukh et al. 2022; Lu et al. 2022), but it often involves only relatively subtle disruptions to key gametogenic processes, leaving the broader reproductive machinery intact. A common example in asexual hybrids is the distortion of the cell cycle towards premeiotic genome endoreplication (PMER) (Dedukh et al. 2020), Fig. S1, where the entire genome duplicates before meiosis. This allows for the formation of bivalents between identical chromosomes, effectively preventing genetic variation among progeny despite the occurrence of normal crossovers. It also circumvents the need for homologous chromosome pairing from divergent parental species, which may fail to align properly.

The emergence of hybrid asexuality is frequent in some taxa like fishes, amphibians, and reptiles (Stöck et al. 2021). Experimental crossings of loaches (Cobitidae) have shown that in eight out of eleven randomly formed species pairs, asexuality was the sole reproductive outcome, demonstrating PMER as a dominant mechanism (Marta et al. 2023). This underscores a complex relationship between hybrid sterility and asexuality, suggesting that hybrid asexuality may be a crucial phase in the speciation process (Janko et al. 2018; Reifová et al. 2023), substantially extending the temporal window during which diverging species can produce fertile hybrid lineages, even though parental chromosomes are too incompatible to form proper bivalents (Dedukh et al. 2020).

Thus, asexual hybrids act as ‘evolutionary capsules’, preserving clonal copies of their sexual ancestors’ mitochondrial or nuclear genomes over extended periods, and in extreme cases, such as with *Squalius* and *Hypseleotris* fish and *Ambystoma* salamander hybrids (Alves et al. 2001; Bi and Bogart 2010; Unmack et al. 2019), this preservation extends beyond the extinction of one parental species, leaving its genetic legacy carried solely by its asexual descendants.

For this study, it is crucial to note that although asexuality is often equated with clonality, the genomes within asexual lineages are not static but evolve beyond mere mutation accumulation. Asexual lineages, including those that bypass the first meiotic division or employ PMER, accumulate structural variants, (e.g. Triantaphyllou 1981; Normark 1999) and experience gene conversions that result in Loss of Heterozygosity (LOH) (Tucker et al. 2013; Warren et al. 2018; Janko et al. 2021; Jaron et al. 2021). These genetic alterations may be adaptive, potentially mitigating deleterious mutations (Kočí et al. 2020) and optimizing gene expression (Smukowski Heil et al. 2017; Bartoš et al. 2019; Janko et al. 2021), but importantly, they facilitate allele exchange between the original parental genomes ‘frozen’ since the origin of lineage. Contrary to the conventional view that asexual hybrids are evolutionary dead ends, it also appears that hybrid asexual lineages may not be a terminal fate for their genomes because some lineages maintain ‘clonality’, like gynogenesis, at one ploidy level (typically diploid) but can switch to ‘hemiclonality’ or even meiotic hybridogenesis when ploidy changes, typically when becoming triploid (e.g. Vrijenhoek 1979; Alves et al. 2001; Angers and Schlosser 2007; Dedukh et al. 2024). This flexibility in reproductive modes can occasionally reconnect these lineages to the sexual gene pool. For instance, when a hybridogenetic triploid with an AAB nuclear genome and B-type mitochondria produces haploid A-type eggs, subsequent fertilization by A sperm can produce AA zygotes with B mitochondria. This results in cytonuclear mosaicism, reintroducing asexual genomes into the A species’ gene pool and facilitating mtDNA introgression without corresponding nuclear gene flow (Kwan et al. 2019).

The accumulating evidence of such clonality imperfections and the frequent shifts between clonal and hybridogenetic reproductive modes in natural hybrids raises a crucial question: Could the interplay of these processes also have a significant impact on the gene pool of their sexual parental species, leading to atypical introgressions pathways in nuclear compartments?

*Cobitis* loaches emerge as a particularly suitable model to examine the propensity of asexual organisms to mediate gene flow between asexual and sexual lineages and among parental taxa themselves. Within Europe and Anatolia, loaches form several deeply diverged speciose lineages, like *Cobitis* sensu stricto mostly from non-Mediterranean Europe and the Adriatic and Bicanestrinia lineages from Balkanian and Anatolian rivers (Fig. 1b). Hybridization of *Cobitis* species within and sometimes between such lineages have independently given rise to asexuality on multiple occasions (Janko et al. 2012), and experimental evidence confirmed that hybridization triggers PMER in hybrids of various species pairs (Marta et al. 2023). Despite PMER, some interchange among orthologous chromosomes exists in clones, leading to the selective accumulation of LOH events (Janko et al. 2021). Moreover, some of these lineages have been observed to switch to hybridogenesis and premeiotic genome elimination in response to changes in ploidy (Dedukh et al. 2024).

**Figure 1.**
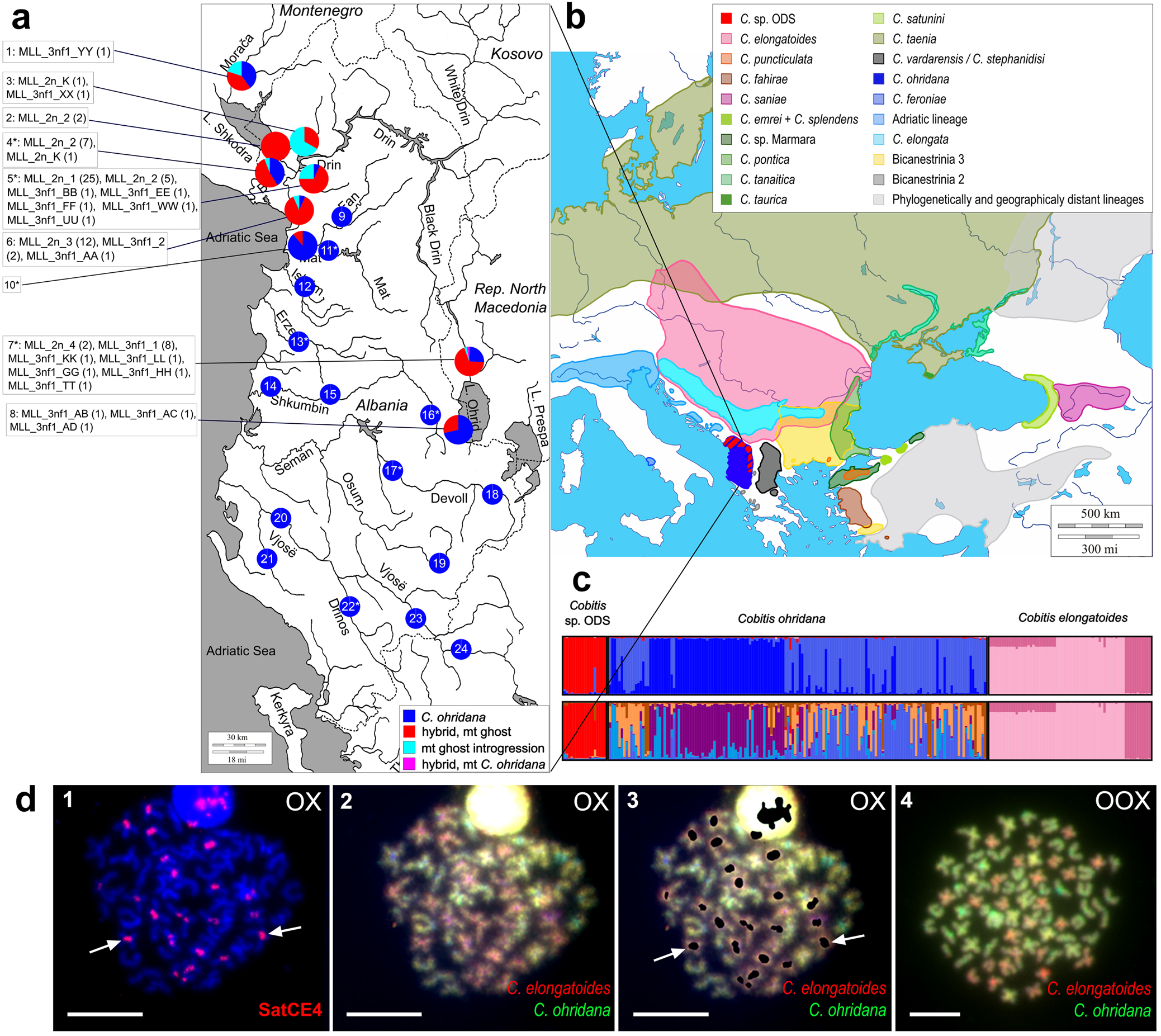
a) Sampling localities and distribution of investigated genomotypes of *Cobitis* in Albania. Pie charts show relative frequency of genomotypes at each locality. For *Cobitis* sp. ODS, multilocus lineages (MLL) and their absolute frequencies (in parentheses) are shown for each locality. Numbers of localities from where samples used for sequence capture originate are marked with asterisk. Locality numbers correspond to those in Table S1. b) Distribution of known *Cobitis* species in Europe and Middle East. Regions where *Cobitis* does not occur in white. Colors correspond to clade colors in Fig. 2. c) Results of Structure analysis with k=5 (upper row) and k=8 (lower row), demonstrating the optimal clustering of diploid individuals sampled in Albania, contrasted against the data from the same loci in *C. elongatoides* as of Janko et al. (2018). (*Note that for Cobitis sp. ODS, we intentionally input only single representative for MLGs in order to avoid an effect of clonal genotype multiplication on Structure’s performance.*) d) Identification of chromosomes of *C. ohridana* and the ‘ghost’ species on somatic metaphase plate from diploid (1-3) and triploid (4) hybrid *Cobitis* sp. ODS individuals. 1) mapping of satCE04 probe (indicated by arrows) which localizes only in centromeric regions of chromosomes from ‘ghost’ species but not in *C. ohridana* chromosomes in diploid hybrid. 2) The same metaphase plate as in (1) after comparative genome hybridization revealing the origin of individual chromosomes (red color chromosomes correspond to *C. elongatoides* and green ones to *C. ohridana*). 3) Merged figure showed the presence of pericentromeric repeat satCE04 probe (indicated by arrows) only in ‘ghost’ genome detected by comparative genome hybridization. 4) Metaphase plate from triploid hybrid showing 50 *C. ohridana* chromosomes and 25 chromosomes corresponding to ‘ghost’ species. Scale bar in d = 10 µm.

In this study, we integrate Europe- and Middle East-wide sampling with cytogenetic, phylogenetic, population genetic, and genomic approaches to investigate the interactions between sexual species and their presumed asexual hybrid descendant. We provide evidence that hybridization between Adriatic species *Cobitis ohridana* and a different, now-extinct, species led to the emergence of a hybrid taxon inhabiting Ohrid-Drin-Skadar (ODS) system in the south-Adriatic biogeographic region (Perdices et al. 2008), tentatively named *Cobitis* sp. ODS. It preserves the parental genome beyond its extinction through clonal passaging but limited recombination between both genomic copies in hybrids induces LOHs. When polyploidized, such hybrids introduce the extinct species’ alleles back into the *C. ohridana* population through hybridogenesis, thereby facilitating a ‘zombie’ gene flow from a long-extinct taxon into an existing sexual species.

## II. RESULTS

### II.a. Phylogenetic results support the co-occurrence of two Cobitis forms across south-Adriatic biogeographic region

Phylogenetic relationships among investigated individuals were evaluated on three nuclear and one mitochondrial markers in Europe- and Middle East-wide context (Figs 1b, 2a-c, e, Table S1-S3). Despite some polytomies in nuclear markers’ phylogenetic trees, all but the *RAG1* indicated *C. ohridana* as the closest relative to Tyrrhenian *C. feroniae*, sometimes sharing alleles. The most variable among nuclear markers, *S7* intron, revealed 32 alleles in *C. ohridana*, divided into two clades diverged from each other by 1.6% p-distance. One is widespread throughout south-Adriatic region, for simplicity further on referred as Albania, appearing as sister to *C. feroniae*, and the other localized to central to northern Albania. In addition, in the ODS system (predominantly in northern and eastern Albania), we found specimens with high degree of heterozygosity in the three nuclear markers (Figs 1a, 2a, b, e), which always harbored one allele shared with *C. ohridana* and another one diverging by 3.1% - 3.3%, that belonged to the *Cobitis* sensu stricto clade, but did not belong to any known extant species of the clade. Such individuals were subsequently called *Cobitis* sp. ODS. In *S7*, the highest similarities of their ‘ghost’ alleles were found to *C. saniae* (p-distance 1.4%), *C. puncticulata* (1.5%), *Cobitis* sp. Marmara (1.8%), and *C. elongatoides*, *C. pontica* and *C. satunini* (2.1%), Figs 1b, 2a, b, e. mtDNA lineages found in *C. ohridana* and *Cobitis* sp. ODS differed by 12.2%. Lineage corresponding to *C. ohridana* contained 41 haplotypes split into geographically distinct sublineages, occupying southern and northern parts of Albania, respectively, with p-distance divergence of 1.3%. *Cobitis* sp. ODS lineage contained 8 haplotypes and clustered within the *Cobitis* sensu stricto clade forming a polytomy with Danubian-Caspian *C. elongatoides/tanaitica/pontica* complex and *C. puncticulata* distributed along the Marmara Sea, differing by p-distance of 2.45% and 2.51%, respectively (Figs 1b, 2c). Evidence of mtDNA exchange between the two lineages was observed, with one *Cobitis* sp. ODS individual harboring a *C. ohridana* mtDNA haplotype and 19 *C. ohridana* specimens from ODS showing mtDNA haplotypes common to *Cobitis* sp. ODS (Figs 1a, 2d). Isolation with Migration analysis confirmed significant mtDNA gene flow between *C. ohridana* and *Cobitis* sp. ODS with the Likelihood Ratio Tests (LRT) rejecting the models of no-migration (2LLR = 1724, p < 0.01) and of unilateral migration from either side (2LLR = 11.96 and 1365, p < 10^-4^). Yet, the model assuming equal migration rates between populations could not be rejected in favor of asymmetric migration (2LLR = 1.257, p = 0.262; Fig. 2d).

**Figure 2.**
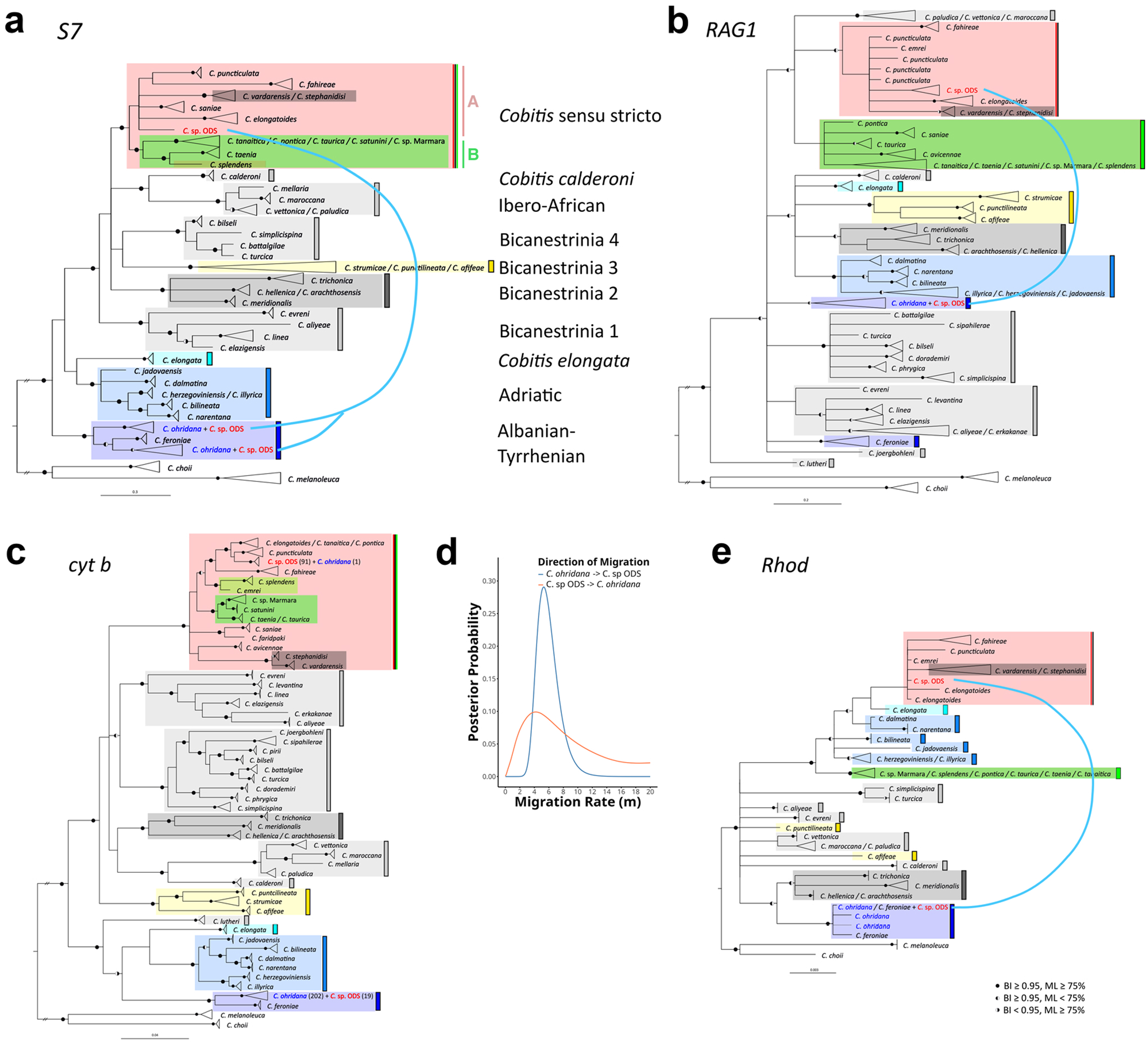
Phylogenetic trees reconstructed by BI phylogenetic reconstruction for three nuclear markers: intron of S7 recombination protein gene (*S7*) (a), exons of Recombination Activating Gene 1 (*RAG1*) (b) and Rhodopsin (*Rhod*) (e), and mitochondrial cytochrome *b* gene (*cyt b*) (c). *Cobitis elongatoides* clade – A, and *C. taenia* clade - B of *Cobitis* sensu stricto lineage shown in (a). Support of nodes (posterior probability, pp, and bootstrap support, bs) given by filled half-circles when pp ≥ 0.95 or bs ≥ 75 %. Coloration of species/lineages follows colors of distribution ranges Fig. 1b. Blue line shows phylogenetic placement of alleles present in *Cobitis* sp. ODS hybrids. *Cobitis* sp. ODS shown in threes in red, *C. ohridana* in blue. d) Summarization of the posterior probability distributions for migration rates between *C. ohridana* and *Cobitis* sp. ODS based on *cyt b* data as obtained from IMa3 software (x axis parameterizes the migration rate; y axis shows the posterior probability).

Thus, the investigated south-Adriatic biogeographic region appears thoroughly populated by *C. ohridana* and by *Cobitis* sp. ODS distributed predominantly in northern and eastern Albania (Figs 1a-b), whose admixed nuclei combines alleles from *C. ohridana* with those of unknown ancestor belonging to the *Cobitis* sensu stricto lineage (Figs 2a, b, e). The relaxed molecular clock calibrated by vicariance along the Gibraltar suggested the divergence between both putative parental lineages at 22.8 - 47.7 Mya in *cyt b* and at 12.7 - 61.5 Mya in *S7* genes, and indicated that ‘ghost’ alleles in *Cobitis* sp. ODS diverged from their nearest existing *Cobitis* sensu stricto counterparts about 1.9 - 6.5 Mya in *cyt b* and 3.1 - 35.8 Mya in *S7* (Fig. S2). While both estimates were subject to large confidence intervals, especially for the less variable *S7* dataset, they clearly suggested early Pleistocene or older divergence of the putative parent of *Cobitis* sp. ODS from other *Cobitis* sensu stricto species.

### II.b. Karyotypes indicate presence of C. ohridana and diploid - triploid hybrid taxon Cobitis sp. ODS

*Cobitis ohridana* had a diploid chromosome number 2n = 50, fundamental number of chromosome arms (NF) = 66 and karyotype composed of 16 meta-/submetacentric (M/SM) + 34 subtelo-/acrocentric (ST/A) chromosomes (Figs S3a-b). *Cobitis* sp. ODS occurred in two karyoforms, one diploid and one triploid. The diploid karyoform contained 2n = 50 chromosomes (NF = 80) with one haploid set of chromosomes apparently resembling the karyotype of *C. ohridana* (8 M/SM + 17 ST/A), and the second set with 22 M/SM + 3 ST/A chromosomes, similar to *C. elongatoides* from the *Cobitis* sensu stricto clade, see Figs S3c-e (Marta et al. 2020). The triploid form (3n = 75, NF = 106) possessed a diploid chromosome complement reassembling *C. ohridana* (16 M/SM + 34 ST/A) and a haploid set with 22 M/SM + 3 ST/A chromosomes again similar to *C. elongatoides* (Fig. 1d). For simplicity we designated *C. ohridana* individuals as ‘OO’, *Cobitis* sp. ODS as ‘XX’, diploid hybrid *Cobitis* sp. ODS individuals as ‘OX’, triploid hybrid *Cobitis* sp. ODS individuals as ‘OOX’.

Ancestral chromosomal sets in diploid and triploid *Cobitis* sp. ODS specimens were distinguished by comparative genome hybridisation (CGH), with the ‘ghost’ genome showing signals after the labelling with a genomic probe of *C. elongatoides* DNA and the other copy, or two copies in triploids, showing signals of probe from *C. ohridana* genomic DNA (Fig. 1d). Pericentromeric satCE04 probe developed from *C. elongatoides* (Marta et al. 2020) further clearly marked to pericentromeric regions ‘ghost’ genome chromosomes in *Cobitis* sp. ODS, while *C. ohridana* specimens used for control did not show any accumulation of such a signal (Fig. 1d). No detectable intergenomic exchanges have been noted between the parental chromosomal sets, suggesting large-scale genomic integrity and an ‘F1’ hybrid constitution of *Cobitis* sp. ODS.

### II.c. Meiotic analysis and reproductive experiments suggest sexual reproduction in C. ohridana, clonal reproduction of diploid Cobitis sp. ODS individuals and meiotic hybridogenesis with occasional cloning in triploid Cobitis sp. ODS

Pachytene as well as diplotene gonocytes of *C. ohridana* males and females possessed 25 well-formed bivalents, each containing at least one mismatch repair protein MLH1 focus (Figs 3a, b, h). Pachytene oocytes of diploid *Cobitis* sp. ODS contained 50 abnormally paired chromosomes. Only a few fully formed bivalents had stained central components of the synaptonemal complex SYCP1, while other chromosomes had partial synapsis or existed as univalents with only lateral SYCP3 components staining. By contrast, diplotenic oocytes contained exclusively 50 bivalents (i.e. 100 chromosomes), of which 25 showed FISH signal of the satCE04 pericentromeric probe, corresponding to the ‘ghost’ genome, and the remaining 25 *C. ohridana* bivalents were without the signal. The presence of oocytes with 50 bivalents suggests their emergence through premeiotic genome endoreplication during gametogenesis of diploid hybrids. The difference between pachytene and diplotene stages is consistent with other *Cobitis* hybrids (Dedukh et al. 2020, 2021; Marta et al. 2023) and suggests that only endoreplicated oocytes can progress to the diplotene stage (Fig. 3c-d, i).

**Figure 3.**
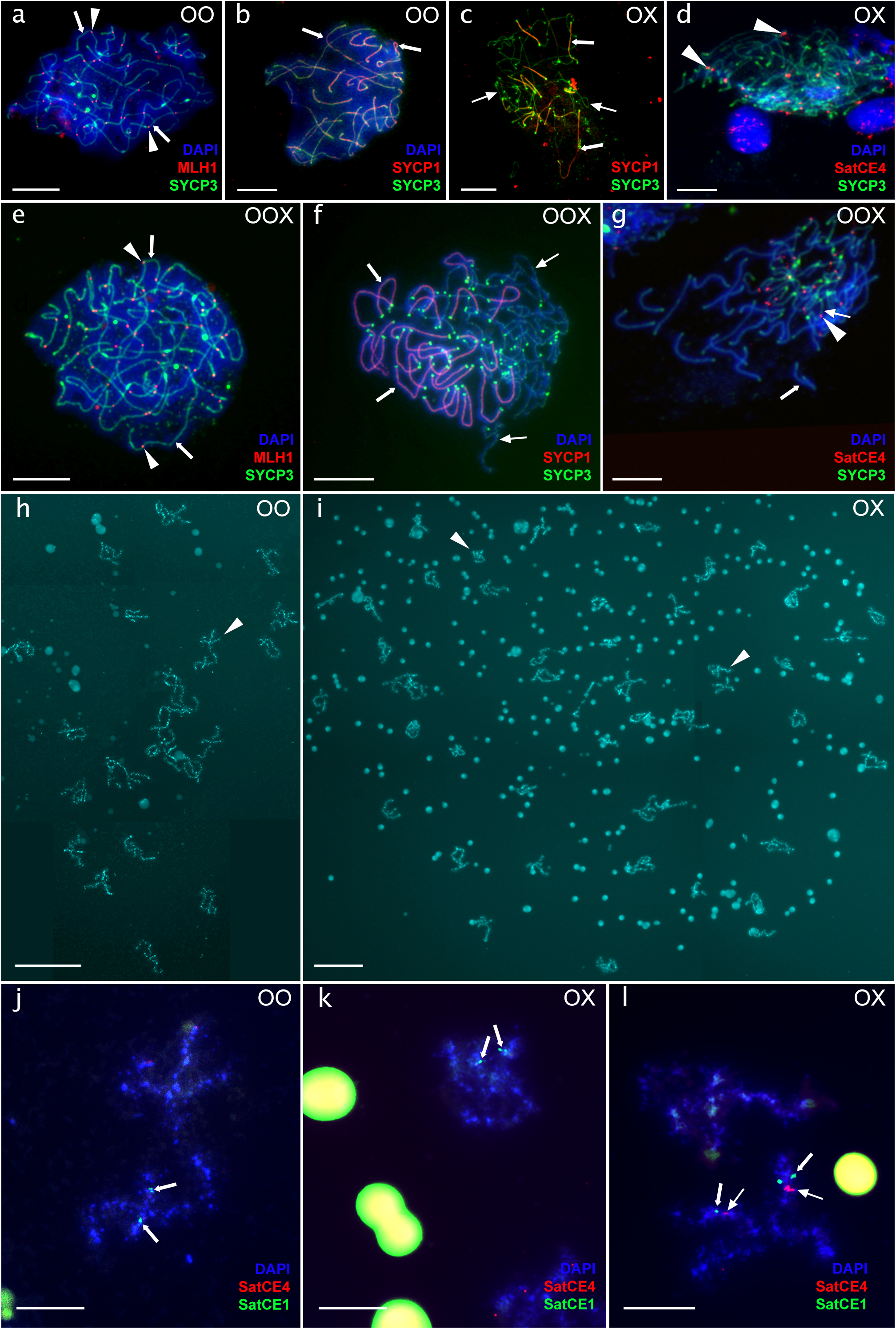
Meiotic spreads during pachytene (a-g) and diplotene (h-i) stages in *C. ohridana* females (a, b, h), diploid *Cobitis* sp. ODS hybrids (c, d, i) and triploid *Cobitis* sp. ODS hybrids (e-g). Thick arrows indicate bivalents, thin arrows indicate univalents, and arrowheads indicate crossing-overs in (a-g). Bivalent formation in *C. ohridana* was confirmed by the presence of crossing-over loci (red staining in a) and the accumulation of central (SYCP1 protein, red staining in b) and lateral (SYCP3 protein, green staining in a, b) components of synaptonemal complexes. The aberrant pairing between orthologous chromosomes was detected in nonduplicated pachytene oocyte (c) from diploid hybrids after immunostaining with SYCP1 and SYCP3 proteins. Centromeric regions on chromosomes corresponding to the ‘ghost’ genome in nonduplicated pachytene oocytes were visualized by FISH with satCE04 repeat (d). Two populations of pachytene oocytes were observed in triploid hybrids: one with 25 bivalents (e), visualized by the presence of crossing overs, and another with 25 bivalents and 25 univalents (f), detected by the accumulation of SYCP3 and SYCP1 proteins. FISH mapping of satCE04 repeat on pachytene oocytes with 25 bivalents and 25 univalents showed signal accumulation only on univalents corresponding to the ‘ghost’ genome (g). Diplotene oocytes of *C. ohridana* (h) contained 25 bivalents with arrowheads indicating a bivalent that is enlarged in (j). This bivalent was detected by the chromosome-specific probe (SatCE1, indicated by thick arrows) and showed a weak signal from the centromeric (SatCE4) probe. Diplotene oocytes of diploid hybrid (i) contained 50 bivalents, consisting of 25 *C. ohridana* bivalents and 25 bivalents corresponding to the ‘ghost’ genome. Arrowheads in (i) indicate the enlarged bivalents in (k, l) which were visualized by chromosome-specific (SatCE1, shown by thick arrows) and centromeric (SatCE4, shown by thin arrows) probes. Bivalent corresponding to *C. ohridana* chromosome (k) showed weak accumulation of centromeric repeat (SatCE4), while the bivalent formed by chromosomes corresponding to ‘ghost’ genome (l) exhibited much intensive accumulation of centromeric repeat (SatCE4, indicated by thin arrows). Scale bars = 10 µm in a-g, and in j-l; scale bars in h-i = 50 µm.

*Cobitis* sp. ODS triploids possessed two types of pachytene oocytes. The first type contained 25 ‘ghost’ ancestor’s chromosomes in univalents with the positive FISH signal from satCE04 pericentromeric probe, in addition to 25 *C. ohridana*-type bivalents without the signal, which demonstrated loaded SYCP1 protein complex. The second type contained only 25 fully formed bivalents with no satCE04 signal, likely containing only *C. ohridana* chromosomes (Fig. 3e-g).

### II.d. Population genetics of mixed populations indicates variable proportion of sexuals and asexuals and moderate levels of clonal diversity

Microsatellite analysis involved 253 diploid and 28 triploid individuals collected from 19 populations across Albania (Table S1). Diversity of diploids (177 individuals without missing data; COTA_006 is not amplified in *C. ohridana*), complemented with 68 *C. elongatoides* samples from Janko et al. (2012) were best explained by k=5 and k= 8 clusters with the largest delta K according to Structure (Fig. 1c). These observations always separated *C. elongatoides*, split into two clusters, from all remaining Albanian samples that were split into 2 or 5 clusters, respectively, and corresponded to *C. ohridana* according to genetic data. An additional single distinct cluster was formed by individuals assigned to *Cobitis* sp. ODS. These individuals always obviously possessed one set of microsatellite alleles shared with *C. ohridana* or similar in size to its alleles, and another set from a different, unknown species. Triploids were not analyzed by Structure software, but clearly possessed in all loci one or two alleles corresponding to *C. ohridana* and the other one to an unknown species, similar to that present in diploid *Cobitis* sp. ODS.

All *C. ohridana* and all triploid *Cobitis* sp. ODS specimens possessed a unique combination of alleles at nine scored loci (Table S4). By contrast, the 57 *Cobitis* sp. ODS diploids could be separated into 18 multilocus genotypes (MLGs), of which the most common involved 9 individuals. Further, analysis of pairwise distances indicated that all 57 diploid *Cobitis* sp. ODS individuals may be grouped into only four clones (Multilocus Lineage; MLL; Tables S1, S4, Fig. 1a), with the most common clone containing 27 individuals and the least dominant two. Among the 28 triploids, the clustering method identified two or three MLLs (clones), depending on the method, which comprised 16 or 10 individuals, respectively, while all remaining triploids appeared independent of the rest (Table S1, S4).

### II.e. Genome-wide ancestry indicates loss of heterozygosity accumulating in hybrids’ genomes

We complemented previously published exome-capture data of *Cobitis* sensu stricto loaches with sexual species from the Balkans and Anatolia and newly sequenced 9 samples of *C. ohridana*, three diploid, and two triploid *Cobitis* sp. ODS hybrids covering major Albanian river systems (Fig. 1a, Table S1). Mapping the reads against the published *C. taenia* reference transcriptome (Janko et al. 2018) retrieved over 400 k quality-filtered bi-allelic SNPs, with *Cobitis* sp. ODS individuals having significantly higher genome-wide heterozygosity than any sexual species used in this study (t-test p = 1.63e-05).

Single nucleotide polymorphisms (SNPs) of diploid and triploid *Cobitis* sp. ODS hybrids were categorized according to Ament-Velásquez et al. (2016) and Janko et al. (2021) relative to their putative parental lineages, one being represented by *C. ohridana* and the other, the ‘ghost ancestor’ by a pool of all *Cobitis* sensu stricto species. In each hybrid we detected between ∼ 8k and ∼ 9k so-called ‘private-hybrid’ SNPs (i.e. categories private_hybrid 1a and 1b in Table S5), not found in any parental species, potentially representing new mutations in clonal lineages or those inherited from an extinct ancestor (Ament-Velásquez et al. 2016; Janko et al. 2021). Of these, 99% were heterozygous, aligning with expectations for asexual reproduction, when new SNP variants are maintained on that ancestral subgenome, where they appeared originally.

Of the 27,311 so-called diagnostic SNPs differentiating all *C. ohridana* specimens from all *Cobitis* sensu stricto species (categories shared 3a, 3b11 and 3b12 in Table S5), majority were heterozygous in *Cobitis* sp. ODS hybrids. However, between 4%-5.8% such positions exhibited LOH, with a significant bias towards preserving *Cobitis* sensu stricto alleles (63%-65%) in diploids, whereas triploids showed a higher retention of *C. ohridana* alleles (∼62%); Table S5.

The normalized coverage patterns at LOH sites were multimodal, with a dominant peak around 1, indicative the gene conversions preserving chromosomal copy numbers, and minor peaks at coverages ∼ 0.5 in diploids and ∼0.33 and ∼ 0.66 in triploids, respectively, suggesting events of allelic losses (Tucker et al. 2013; Janko et al. 2021). A nonlinear least squares approach using gamma distributions confirmed diploid datasets were best fitted by a mixed two-gamma distribution, representing gene conversions occurring at ∼65% - 76% LOH sites and hemizygous deletions on remaining ones (Table S6, Figs 4a-b). Triploids were generally best fitted by a more complex three-gamma model, incorporating gene conversion occurring at ∼ 34% - 50% LOH sites, with hemizygous and double deletions on remaining ones, but LOH sites preserving *C. ohridana* allele were best fitted by a mixed two-gamma distribution without double deletions (Table S6, Figs 4c-d).

**Figure 4.**
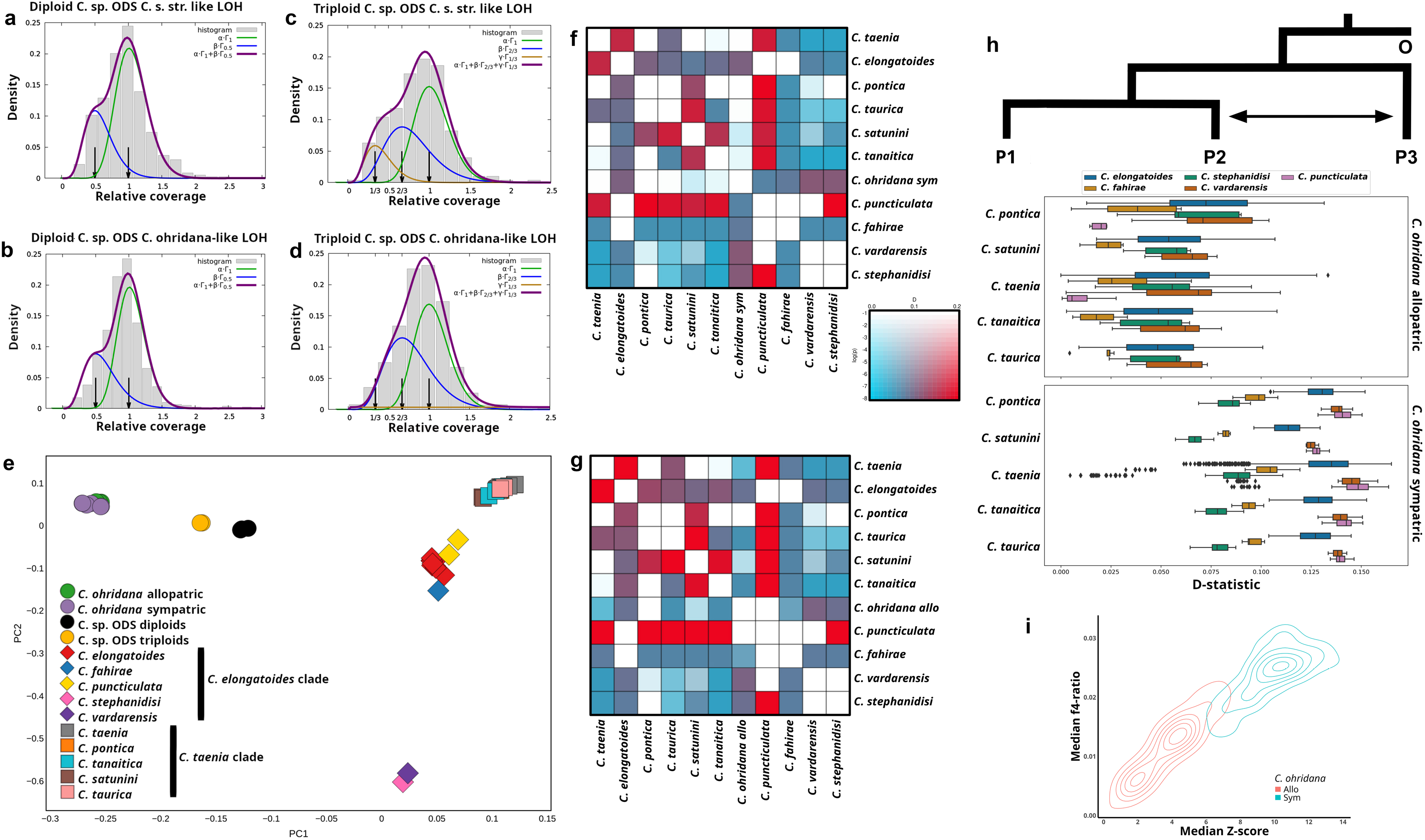
a-d) Histograms of relative coverages at LOH loci in representative diploid (a-b) and triploid (c-d) *Cobitis* sp. ODS hybrids. Arrows depict biologically meaningful values for given ploidy, i.e. 0.5 and 1 for diploids and 0.33, 0.66 and 1 for triploids; green lines show the fit of a single Gamma distribution with mean centered at value 1 representing gene conversions, blue lines show the fit of a single Gamma distribution centered at 0.5 or 0.66 representing hemizygous deletion for diploids and triploids, respectively, and brown lines show the fit of a single Gamma distribution centered at 0.33, representing double deletions in triploids. Thick purple lines represent the best fit mixture of two or three Gamma distributions with the coefficients α, β (and γ) indicating the proportions of each Gamma distribution in the combined model. e) Multidimensional scaling of SNP variability among exome-capture data from sexual species. f-g) Heat matrix demonstrating the detected admixture through D values and corresponding p-values for all species combinations, using *C. ohridana* populations sympatric (f) and allopatric (g) with hybrids, respectively. h) Triplet test of introgression between *C. ohridana* populations sympatric or allopatric with hybrids *Cobitis* sp. ODS, respectively, and *Cobitis* sensu stricto. D values of ABBA-BABA are provided for every combination of *C. ohridana* specimens positioned at P3 position, and *C. elongatoides* clade of *Cobitis* sensu stricto lineage positioned at P2 and *C. taenia* clade of *C.* sensu stricto lineage positioned at P1. The arrow indicates gene flow between species at P2 and P3 positions. i) 2D plot of median Z scores and corresponding f4 values measured for all combinations of *Cobitis* sensu stricto and *C. ohridana* presented in (h).

### II.f. Historical admixture between C. ohridana and Cobitis sensu stricto species

The distribution of SNP variant among samples revealed significant genetic divergence among *Cobitis* sensu stricto, Adriatic, and Bicanestrinia lineages and moderate intraspecific differentiation between *C. ohridana* samples from the ODS river system and those from central and southern Albania (Fig. 4e).

The D statistics, Z scores, and f4 ratios calculated by the Dsuite toolkit indicated possible gene flow between various species pairs belonging to both major groups of *Cobitis* sensu stricto, i.e. between the *C. taenia* clade (*C. taenia*, *C. taurica*, *C. tanaitica*, or *C. pontica*) and the *C. elongatoides* clade (*C. elongatoides*, *C. puncticulata*, *C. vardarensis*, *C. stephanidisi* or *C. fahirae*) (Figs 4f-g), which is consistent with previous findings by Janko et al. (2018). Notably, the introgression was also suggested to occur between *C. ohridana* and some *Cobitis* sensu stricto species. Specifically, the software assumes a ((P1,P2),P3),P4) species tree topology and tests for introgression between P2 and P3 taxon. When *C. ohridana* individuals were placed on the tree branch P3, with any species from the *Cobitis taenia* group positioned at P1, and any species of the *C. elongatoides* group at P2, there was a notable excess of (P1,(P2,P3)),P4 site patterns over (P2,(P1,P3)),P4 site patterns. This resulted in elevated D values and f4 ratios and Z scores, which exceeded 3 when P1 was occupied by either *C. tanaitica* or *C. pontica*, underscoring the statistical significance of the introgression detected (Fig. 4h), indicating introgression between *C. ohridana* and the *C. elongatoides* clade.

Interestingly, these introgression rates appeared higher when using *C. ohridana* specimens on P3 position that came from localities sympatric with *Cobitis* sp. ODS hybrids (Figs 4f-g). This resulted in higher D values, Z scores, and f4 ratios, compared to those from allopatric ones (Figs 4h-i). This difference was significant with GLMMs controlling for random variations among individuals in the P3 position suggested increase of 0.013538 (SE = 0.003104, df = 9.018904, t = 4.362, p = 0.00181) for D values, 0.8141 (SE = 0.2134, df = 9.0169, t = 3.816, p = 0.0041) for Z scores and 0.0039229 (SE = 0.0009261, df = 9.0158663, t = 4.236, p = 0.00218) for f4 ratios. Variance estimates were low across all models with random intercepts variance of 1.783e-05 (Std.Dev = 0.004222) for D values, 0.08581 (Std.Dev = 0.2929) for Z scores, and 1.597e-06 (Std.Dev = 0.001264) for f4 ratios. A block design examining the differences between sympatric and allopatric groups across various genetic backgrounds corroborated the significantly higher introgression rates into *C. ohridana* specimens sympatric with *Cobitis* sp. ODS hybrids than into allopatric ones (sum of ranks W = 20741 with p-value = 3.88 × 10^-12 for D statistic, W = 21173 with p-value of 1.19 × 10^-13 for Z scores and W=20863 with p-value=1.48 × 10^-12 for f4 ratios).

## III. DISCUSSION

### III.a. Evolutionary time capsules: preserving extinct genomes through asexual hybrids

Hybrid forms are known to impact on evolution in various ways but European loaches highlight another unique role: the preservation of extinct genomes through asexual reproduction of hybrid lineages. Rivers of south-Adriatic biogeographic region appear populated by *C. ohridana*, alongside a hybrid form denoted as *Cobitis* sp. ODS occurring in both diploid and triploid states. It combines *C. ohridana* alleles with those phylogenetically belonging to *Cobitis* sensu stricto clade but with no match to any current European or Anatolian *Cobitis* species. Despite extensive sampling, the closest matches are with alleles from Danubian *C. elongatoides* and western Anatolian *C. puncticulata*, with an estimated divergence of approximately 2 million years ago, indicating that *Cobitis* sp. ODS hybrids may be conserving a genome from a species that is now likely extinct, in a form of stable and widespread hybrid lineage.

Cytogenetics indicated that *Cobitis* sp. ODS preserves and transmits both parental chromosome sets without remarkable rearrangements, consistent with the vast majority of SNPs diagnosing *C. ohridana* from *Cobitis* sensu stricto being maintained in a heterozygous state. Such a ‘frozen F1’ state, to a large extent, is unlikely through standard sexual reproduction but instead indicates some form of asexuality. Indeed, diploid and triploid *Cobitis* sp. ODS formed several distinct multilocus lineages (MLLs), which may be considered clones, and diploid *Cobitis* sp. ODS females possess most oocytes with aberrant pairing of chromosomes from both parental haploid sets but also a minor population with a doubled chromosome number forming 50 bivalents instead of 25 Figs 5b, S1. This aligns with PMER, a common gametogenesis alteration among asexual vertebrates and specifically among Cobitidae, which typically occurs in the terminal stage of a minor proportion of oogonial cell development before meiosis, thus enabling clonal reproduction (Lutes et al. 2010; Dedukh et al. 2020, 2021; Marta et al. 2023).

**Figure 5.**
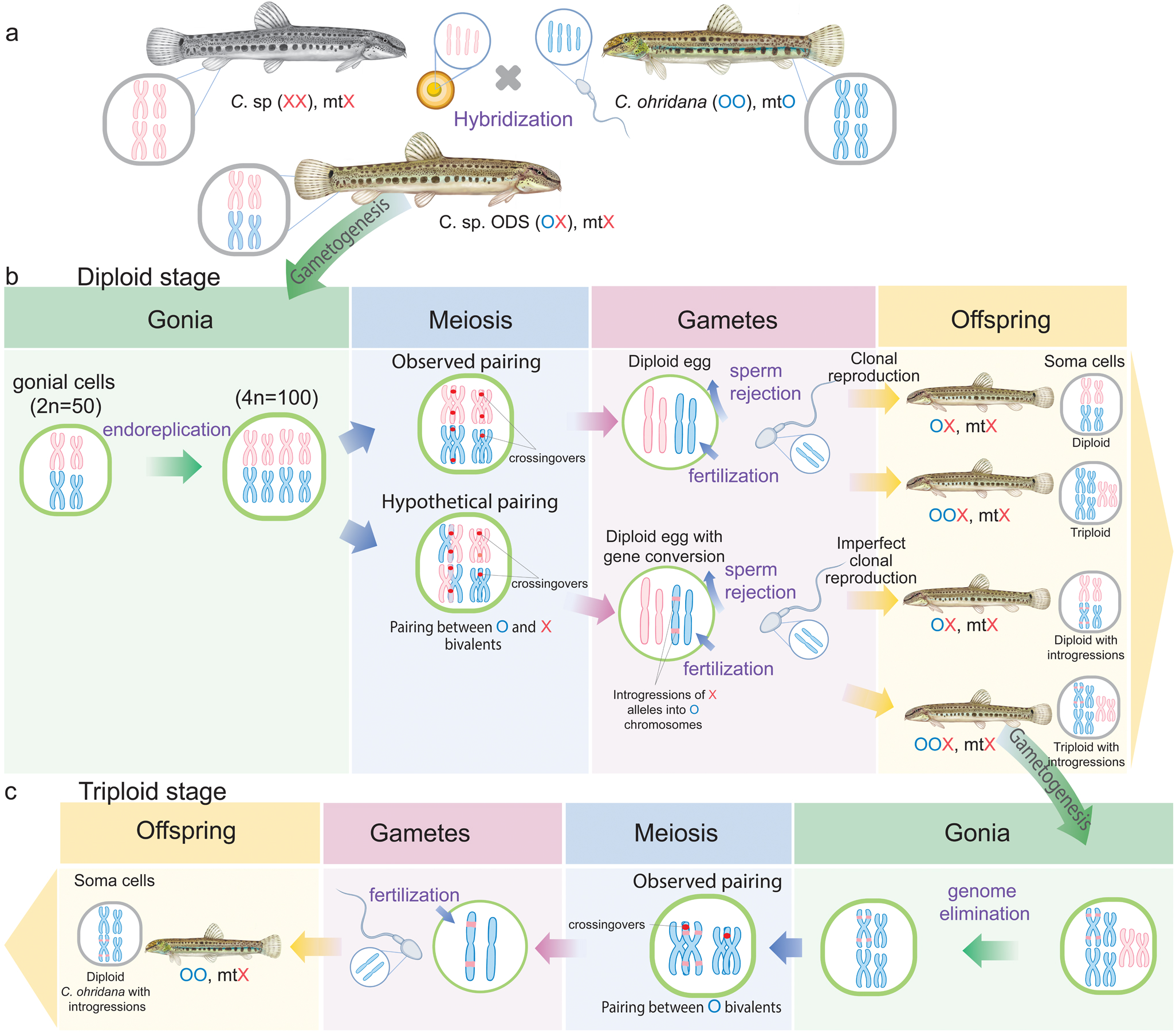
Summary of known and putative reproductive pathways in *Cobitis* sp. ODS hybrid asexuals. For reasons of clarity, this figure shows only gametogenic pathways relevant to the main message of the paper. For schematic description of all observed gametogenic types, please refer to Fig. S1. a) Primary crosses of two parental sexual species, *Cobitis ohridana* (OO, marked in blue) and ‘ghost’ species (XX, marked in pink) resulted in diploid *Cobitis* sp. ODS (OX). Such hybrids include the mitochondrial DNA (designated as ‘mt’) from ‘ghost’ species (X). b) Diploid hybrid females (OX) have premeiotic genome endoreplication, which enables the bivalent formation during the meiotic prophase and recombination between identical chromosome copies (upper panel). Such oocytes resulted in diploid gametes which can be either activated by sperm of *C. ohridana* male causing the emergence of completely clonal diploid progeny. Alternatively, sperm genome can be incorporated into the diploid egg genome leading to ploidy elevation and triploid progeny. In addition, occasional pairing of O and X chromosomes in oocytes with endorepicated genome may lead to gene conversion events (lower panel). Resulted diploid eggs can be also activated by *C. ohridana* sperm leading to diploid clonal hybrids. Additionally, such eggs can incorporate sperm genome leading to the emergence of triploid hybrids. Both diploid and triploid hybrids retain O maternal genome with introgressed X alleles and mitochondrial DNA from ‘ghost’ species (X). c) Triploid *Cobitis* sp. ODS females (OOX) produce several types of gametes. Some oocytes contain only bivalents formed by only *C. ohridana* chromosomes, suggesting the premeiotic elimination of X genome. These oocytes produced haploid gametes, which after fertilization by *C. ohridana* sperm lead to the emergence of *C. ohridana* individuals with introgressed alleles from X genome and mitochondrial DNA from the ‘ghost’ species (X).

Asexual reproduction thus appears to effectively conserve the genetic integrity of the ‘ghost’ ancestral genome across multiple generations, making *Cobitis* sp. ODS hybrids evolutionary living ‘time capsules’, safeguarding genetic blueprints of species that may no longer exist in their pure forms, akin to some other hybrid taxa that may have outlived or replaced their pure-species progenitors (e.g. Alves et al. 2001; Bi and Bogart 2010; Unmack et al. 2019).

### III.b. Biased gene conversion between subgenomes in asexual hybrids

Despite PMER generally ensuring clonal reproduction (Lutes et al. 2010; Majtánová et al. 2016; Janko et al. 2021) Albanian *Cobitis* sp. ODS hybrids accumulated fine-scale genomic restructuring, with approximately 5% of their diagnostic SNP sites exhibiting loss of heterozygosity. These LOH events accumulate through a mix of processes involving allelic deletions and gene conversions, as indicated by multimodal coverage distributions. Peaks at normalized coverages around 0.5 and 0.66 in diploids and triploids respectively suggest hemizygous deletions; a peak at 0.33 in triploids points to double deletions. As shown in other *Cobitis* hybrids (Janko et al. 2021), Albanian triploids also appear more tolerant to allelic deletions, which constitute about 50% of their LOH events, compared to only 25%-35% in diploids. This may suggest that deletions are less impactful on gene product stoichiometry in triploids when altering their genome dosage by approximately 33%, compared to 50% in diploids (Veitia et al. 2013).

Largest part of LOH sites nevertheless possessed normalized coverages around 1 implying gene conversions (Tucker et al. 2013; Janko et al. 2021). These could result from the occasional meiotic pairing of orthologous rather than homoeologous chromosomes following PMER, (Fig. 5b), a process previously hypothesized for other asexual systems too (Janko et al. 2021; Jaron et al. 2021). Alternatively, mitotic conversions — increasingly recognized for their importance in organisms (Jia et al. 2021) — may also contribute to these patterns.

Interestingly, LOH accumulation in diploid hybrids is notably biased towards replacing *ohridana*-like alleles with those from the extinct parental subgenome. Triploids preferentially retained *ohridana*-like alleles, likely reflecting detection limits of our method, where loss of a single *C. ohridana* allele in triploids is masked by remaining heterozygosity. Even so, ∼38% of triploids’ LOH sites still show loss of *C. ohridana* alleles, many conforming to gene conversions rather than deletions, indicating repeated occurrences of gene conversions at homologous sites overwriting both *C. ohridana* alleles. Biased retention of one subgenome, observed in some hybrids and allopolyploids, has been attributed to expression bias when a less expressed subgenome is supposedly replaced or eliminated (e.g. Yoo et al. 2014). However, underlying mechanism is more complex as found in loaches. Notably, non-Mediterranean *C. elongatoides-taenia* and *C. elongatoides-tanaitica* hybrids preferentially retain *C. elongatoides* subgenome despite its overall transcriptional suppression (Bartoš et al. 2019; Janko et al. 2021). Similarly, Albanian hybrids preferentially retain the ‘ghost’ subgenome coming from the *C. elongatodies* clade, suggesting this type of subgenome is consistently preferentially retained over its counterpart even among independently arisen hybrids. The mechanisms underlying observed biased gene conversions remain to be fully elucidated. It could be influenced by uneven distribution of epigenetic modifications, affecting the recombination landscape among parental subgenomes (Mirouze et al. 2012), or by selective pressures favoring homozygosity in certain gene products (Janko et al. 2021).

In any case, our findings document the inherent imperfections in clonal reproduction via PMER, leading to genetic erosion and accumulation of LOH that causes short-range introgression of loci from one subgenome onto the other subgenome’s background (Fig. 5b).

### III.c. Imperfect clonality and ploidy-dependent switches to hybridogenesis facilitate genetic introgression into sexual populations

Genomic mixing within clones may have interesting consequences for coexisting sexual species if asexuals may alter their reproductive modes. In Albanian *Cobitis* sp. ODS hybrids, diploids form widespread natural lineages likely reproducing gynogenetically through PMER, but triploidization through incorporation of *C. ohridana* sperm leads more complex pathways.

On one hand, population genotyping suggests some triploids may also form clonal lineages, potentially even accumulating specific LOH events. However, most triploids display unique multilocus genotypes (MLGs), indicating their de novo origins from diploids. Cytology of triploid females showed selective elimination of the haploid genome from the extinct ancestor, followed by meiotic pairing between the remaining two *C. ohridana* subgenomes, likely leading to recombined haploid eggs. Mechanisms driving such alternation between diverse reproductive strategies remain elusive and may involve factors such as genome dosage, altered sex chromosome dynamics, and epigenetic modifications (Bartoš et al. 2019; Stöck et al. 2021; Lafond and Angers 2024). Nevertheless, they have been reported in other diploid-triploid hybrid fishes (Alves et al. 2001; Dedukh et al. 2024; Lafond and Angers 2024) and amphibians (Dedukh et al. 2015), suggesting they may be common among asexual hybrids, rather than being unique to our system.

Importantly, in combination with imperfect clonality allowing intergenomic gene conversions in diploid hybrids, a switch to meiotic hybridogenesis allows hybrid’s chromosomes to re-enter the sexual population pool, if haploid eggs of a triploid are fertilized by males of the sexual parental species (Fig. 5c). This facilitates the introgression of alleles from extinct species into the sexual host which we have documented on two levels.

First, haploid *C. ohridana*-like oocytes of triploids carry mitochondria from an extinct maternal progenitor, and hence their fertilization by *C. ohridana* sperm produces progeny with diploid sets of *C. ohridana* chromosomes but allochthonous mtDNA. This is consistent with mtDNA introgression observed in the ODS system and mirrors similar findings in other hybridogenetic systems (Alves et al. 2001; Plötner et al. 2008; Kwan et al. 2019), underscoring their evolutionary significance.

For the first time, however, we found that gene conversions of one parental’s alleles onto the other’s chromosome background in diploid asexuals, enables their introgression into the coexisting sexual (*C. ohridana*) population through meiotic triploid hybrids (Figs 5a-c). This process is supported by significantly higher rates of gene exchange in *C. ohridana* populations that are sympatric with the diploid-triploid hybrid *Cobitis* sp. ODS, compared to those in allopatric regions (Figs 4 f-i). Although applied phylogenomic analysis does not specify the direction of introgression, ABBA-BABA analyses indicate that these dynamic interactions lead to approximately 3% of the recipient’s genome in sympatric regions being influenced by introgressive events, implying a significant impact of asexual hybrids on the genetic structure of coexisting sexual populations.

### III.d. Rethinking asexual hybrids in evolutionary context and biodiversity management

Asexual lineages are garnering renewed interest from evolutionary biologists and ecologists, who recognize non-Mendelian reproductive mechanisms as integral to the speciation continuum (Stöck et al. 2021; Reifová et al. 2023). This study highlights an unexpected role of asexual hybrids in efficiently preserving subgenomes of parental species beyond their extinction and reintroducing their genes, transmitted asexually over generations, into the gene pool of contemporary sexual species, thereby mediating sort of ‘zombie gene flow’ (see Fig. 5a-c). Components of this evolutionary mosaic, such as gene conversions, and ploidy-dependent reproductive shifts, have been documented in isolation across various taxa (Alves et al. 2001; Plötner et al. 2008; Tucker et al. 2013; Warren et al. 2018; Janko et al. 2021; Jaron et al. 2021; Dedukh et al. 2024). Thus, prerequisites for processes observed in this study are not exclusive to loaches and analogous processes could be potentially observed in other systems if similar multidisciplinary approaches are applied.

Evolutionary implications of such ‘non-Mendelian’ introgressive hybridization warrant further research focus due to their unique impact on recipient species, differing fundamentally from classical hybrid introgressions. Re-introgression of asexually transmitted chromosomes back to sexual’s gene pool can introduce accumulated deleterious mutations as well as potentially beneficial allelic combinations even enhancing the evolution of ‘super genes’ (Janko et al. 2023). Gene conversions, which mix ancestral subgenomes, are not randomly distributed across the hybrid genome (Janko et al. 2021), suggesting that their introgression into sexual populations can systematically influence specific genomic regions, uniquely shaping their evolutionary trajectories.

Interspecific hybrids are increasingly recognized as significant ecological and evolutionary entities warranting explicit inclusion in conservation strategies (e.g. Jackiw et al. 2015; Chan et al. 2019; Quilodrán et al. 2020; Draper et al. 2021)). However, asexual hybrids remain on the periphery of these discussions, sometimes explicitly excluded from conservation or even taxonomic efforts (rev in e.g. Majeský et al. (2017)). Our findings of dynamic interconnections between sexual and asexual populations underscore an underrated importance of asexuals. In light of current debates on de-extinction, which explore bioengineering methods to resurrect extinct taxa (Shapiro 2017), asexual lineages with their oscillating clonal-hybridogenetic modes and preservation of ancient genomes could provide innovative pathways for recreating extinct parental forms.

In conclusion, our study calls for a more inclusive approach that recognizes the conservation value of asexual hybrids and reconsideration of their legislative status to ensure that these biologically significant organisms are actively integrated into biodiversity management strategies.

## Material and methods

### Samples

518 specimens of spined loaches were obtained from 24 localities throughout the southern-Adriatic biogeographical region (*sensu* Oikonomou et al. (2014)) and used for various downstream analyses (Fig. 1a; Table S1). Additional material of European and Middle East spined loach species (178 specimens) was analyzed for comparative purposes (Table S2).

### Molecular biological techniques

#### Sanger sequencing and cloning of nuclear markers

Total genomic DNA was extracted from fin tissue using Geneaid Genomic DNA Mini Kit (Tissue). We amplified and sequenced three nuclear genes and one mitochondrial gene, namely the first intron of the gene coding for the S7 ribosomal protein (*S7*), Rhodopsin gene (*Rhod*), Recombination Activating Gene (*RAG1*), and the cytochrome *b* gene (c*yt b*), following Marta et al. (2023).

#### Microsatellite analysis

Selected specimens from 19 localities were genotyped using the available cross-species amplifying microsatellite markers run in two multiplexes; multiplex 1: Cota 006, Cota 010, Cota 032, Cota 068, Cota 093, Cota 111, multiplex 2: Cota 033, Cota 037, Cota 041; (see Janko et al. 2012). GeneMarker v. 1.8 (Softgenetics Company) was used to genotype microsatellites data from electropherogram obtained from sequencer.

#### Exome-capture analysis

DNA from 9 *C. ohridana* and 3 diploid and 2 triploid *Cobitis* sp. ODS hybrids covering major Albanian river systems was extracted by standard phenol chloroform protocol, checked by nanodrop(tm) and qubit(tm) (BR 1000 reagents(tm)) instruments and fragmented on Bioruptor(tm) instrument (high mode) by 40 cycles of sonication (30 seconds of cooling in each 60 s cycle). Target enrichment (Seq-Cap) libraries targeting ca 10,000 coding loci were prepared using probes and protocol of Janko et al. (2018). Indexed samples were equimolarly pooled and sequenced in Genecore institute (EMBL, Heidelberg, Germany) in pair-end mode on NextSeq 500 Illumina instrument (medium output).

### Data analysis

#### Phylogenetics

Sanger sequences were verified and edited in Chromas v2.6.4 and aligned using ClustalW within the BioEdit software package v7.2.6.1 (Hall 1999). Heterozygotic nuclear sequences were phased into alleles using Phase v2.1.1 (Stephens et al. 2001) with a probability threshold of 0.9. PCR products of nuclear genes from individuals too heterozygous to be phased were ligated into the pGEM-T vector (Promega), transformation using TOP10F Competent Cells (Invitrogen) and selecting transgenic bacterial clones with blue-white screening. Each clone was amplified and purified using the Wizard Plus SV Minipreps DNA Purification System (Promega) and sequenced using amplification primers. Given the presence of indels in the *S7* dataset, we produced two separate datasets: one containing long sequences from cloned hybrids and sequences without indels, and another trimmed to the first indel. The former was used for phylogenetic reconstruction, while the latter was employed for genotyping. We checked the amino acid translations for stop codons using MEGA11 (Tamura et al. 2021) and ORF Finder (bioinformatics.org), and calculated haplotype diversity and Watterson’s θ using DnaSP v6.12.01 (Rozas et al. 2017).

Single locus phylogenetic trees were reconstructed using Bayesian Inference (BI) and Maximum Likelihood (ML) methods, including previously published sequences of *C. ohridana* as well as of other European and Middle East *Cobitis* species to put our data into broader context (Tables S2-S3).

Unique haplotypes identified using FaBox (Villesen 2007) were used for phylogenetic reconstruction. Best-fitting nucleotide substitution models and partitioning schemes for positions inside codons were selected with PartitionFinder 2 for coding markers and with jModelTest 2.1.7 (Darriba et al. 2012) for the non-coding one. For bayesian phylogeny, MrBayes v3.2.2 (Ronquist et al. 2012) was run with four independent MCMC runs for 2 (*cyt b* and *RAG1*) and 1 million generations (*Rhod* and *S7*) while sampling trees every 2,000 and 1,000 generations, respectively. Convergence was assessed with Tracer v1.7.0, and the first 25% of trees discarded as burn-in, and a 50% majority-rule tree constructed. For ML tree reconstruction, the RAxML 8.2.12 (Stamatakis 2014) was run through the CIPRES Science Gateway portal with node support estimated from 1,000 bootstrap replicates. Uncorrected p-distances were calculated using MEGA 11.

For the two markers with highest variability (*cyt b* and *S7*) we constructed time-calibrated trees with uncorrelated clock model using previously published calibration point corresponding to the opening of the Gibraltar strait (∼5.33 MYA), which separated two sister species, i.e., *C. maroccana* and *C. mellaria* (Doadrio and Perdices 2005). Although the usage of this recent calibration point may have limitations with respect to deeper nodes, it is to date the only relevant calibration point for this group. In order not to violate the interspecific branching prior (Yule model), we selected one individual per species. For *C. ohridana* and ‘ghost’ species, we selected the most frequent haplotype (*cyt b*: O4, X1, respectively; *S7*: O1, X1, respectively). We prepared the analysis in BEAUti v. 1.8.0 with the substitution models from jModelTest v. 2.1.7 (Darriba et al. 2012) and ran in BEAST v. 1.8.4 (Drummond et al. 2012) under the uncorrelated lognormal clock for each marker, which adequacy was verified by the Likelihood Ratio Test (LRT) (Huelsenbeck and Crandall 1997) in MEGA11. Speciation Yule prior was applied with the uniform birth rate (lower=0, upper=100, initial=1) and uniform distributions were set for relative rates of substitution between pairs of nucleotides (lower=0, upper=100, initial=1). The prior for tmrca of clade defined by calibration point was set to normal distribution. The ucdl.means were set to exponential (initial=1, mean=1, offset=0). The remaining priors were set as default. BEAST was run three times with 20 and 30 million generations for *cyt b* and *S7*, respectively, saving parameters and trees every 2,000 and 3,000 generations. Adequate mixing of the MCM chains and ESS values were checked with Tracer v. 1.7.1 (Rambaut et al. 2018), LogCombiner v. 1.8.4 was used for merging tree files with 10% burn-in and TreeAnnotator (Drummond et al. 2012) v. 1.8.4 was used for identifying the maximum clade credibility tree. All trees were visualized in FigTree v. 1.4.4 (Rambaut et al. 2018) and edited in Inkscape v. 0.92.3.

#### Population genetics

We checked the microsatellite data in MICRO-CHECKER V.2.2.3 (Van Oosterhout et al. 2004) for null-alleles, large allele dropouts and scoring errors. Summary statistics, deviations from Hardy-Weinberg equilibrium (HWE), *F*_ST_, and pairwise linkage equilibria were evaluated for each locus per taxon using MSA v. 4.05 (Dieringer and Schlötterer 2003), Genepop v.4.1.3 (Rousset 2008) and FSTAT v.2.9.2 (Goudet et al. 2002). To study structure of population, we used the Structure software v. 2.3.4 (Falush et al. 2003) with following parameters: 10,000 steps of burn-in periods; 10,000 MCMC steps; Ancestry Model – Admixture Model; Allele Frequency Model – Allele Frequencies Correlated. Simulations were run for K from 2 to 6 with 5 iterations each, and best K value selected with the Structure Harvester (Earl and vonHoldt 2011). CLUMPP v. 1.1.2 (Jakobsson and Rosenberg 2007) was used to merge and average data for the best value of K to obtain one final barplot for the Distruct v. 1.1 (Rosenberg 2004).

The GenAlEx 6.5 software (Peakall and Smouse 2012) was used to identify all unique multilocus genotypes (MLG). As described in Janko et al. (2012), we further identified groups of MLGs that are related to each other more closely than expected by chance alone and might therefore represent members of the same clone, the so-called multilocus lineage (MLL). To do so, we calculated the sum of the differences in allele lengths between each pair of MLG (Meirmans and Van Tienderen 2004) and subsequently simulated hybrid genotypes by 10,000 random combinations of individuals from hybridizing species to obtain the null distance distribution assuming each MLG originated from independent sexual intercourse between parental species. Since the parental from the *Cobitis* sensu stricto clade was missing, we used published microsatellite data of *C. elongatoides* (Janko et al. 2012) as a proxy for the extinct parental’s variability. Empirical and simulated distance distributions were compared to evaluate the probability that any two MLG belong to the same clone after correcting for multiple comparisons, with the false discovery rate correction for multiple comparisons.

To evaluate the MLL clustering in triploids, two simulations were run. First, we used analogous assumption of independent intercourses as in diploids and simply randomly combined alleles from two *C. ohridana* and single *C. elongatoides* specimen to obtain random samples of triploid with two *C. ohridana* and one ‘ghost’ genomes. In the second approach, to take into account variability of putatively clonal diploid hybrids (Janko et al. 2012), we each time randomly selected one diploid hybrid as a maternal ancestor and added a haploid set of alleles of randomly selected *C. ohridana* as a paternal one. Both types of simulated distance matrices were again compared to the empirical distance matrix of triploids.

#### *Analysis* of exome-capture data and detection of Loss of Heterozygosity

Raw data were processed using bbduk.sh script from BBmap package v. 39.01 (Bushnell 2014) with the following parameters: *entropy=0.2 entropywindow=50 entropyk=5 ktrim=r k=23 mink=11 hdist=1 ftm=5 tpe tbo ref=adapters.fa* (file provided by BBmap package) *qtrim=rl trimq=10 maq=15 minlen=36*. Bwa mem v. 0.7.17 (Li and Durbin 2009) in default settings was used to align processed reads to *C. taenia* reference transcriptome published and cleaned from potentially paralogous contigs (Janko et al. 2018); https://doi.org/10.5061/dryad.7nt8f). Reads in produced bam files were filtered out based on low mapping quality using samtools v. 1.10 (Danecek et al. 2021) with parameters *-q 10* and *-F 1284*. Variant calling was done using GATK pipeline v. 4.5.0. (Auwera and O’Connor 2020) and genotypic variation among sequenced individuals was assessed using principal component analysis (PCA) based on the resulting single nucleotide polymorphism (SNP) data, filtered with VCFtools (0.1.16) to remove missing values and performed with Plink v1.90b7.2 (Purcell et al. 2007). For each specimen and every contig, we subsequently created a consensus using bcftools consensus v.1.19 with parameters *--iupac-codes --haplotype I* to call heterozygous positions, and inserting ‘N’ in positions with insufficient information using a combination of samtools depth *-aa* and bedtools maskfasta v. 2.29.2. (Quinlan and Hall 2010).

To evaluate the genomic composition of presumed hybrids, we categorized obtained SNPs in exons from nuclear genes into 10 classes based on their distribution among biotypes as outlined by Janko et al. (2021). Assuming that Albanian hybrid taxon *Cobitis* sp. ODS possess genomic copies from two parental species, *C. ohridana* and an unknown species from the *Cobitis* sensu stricto lineage, we compared SNPs against both potential ancestral taxa, using newly obtained as well as previously published studies for ten species of *Cobitis* sensu stricto lineage (Janko et al. 2018, 2021) as a proxy for the ‘ghost’ parental species. Each SNP in the hybrids was classified based on its presence or absence in these potential parental species (Table S5). ‘Species-specific’ SNPs differentiating all *C. ohridana* from all available *C.* sensu stricto specimens (categories shared 3a-b) were crucial for determining whether hybrids were heterozygous, as expected, or homozygous, indicating LOH.

To identify the processes leading to LOH events, we normalized coverages at each LOH site using the ‘total read count’ method (Dillies et al. 2013) and calculated their relative coverages by comparing normalized coverages in hybrids to parental species at the same sites.

Expected relative coverages of ∼1 indicate a process preserving number of allelic copies, indicative of gene conversion, of ∼0.5 or ∼0.66 correspond to hemizygous deletions in diploids and triploids, respectively, and of ∼0.33 to double deletions in triploids (Tucker et al. 2013; Janko et al. 2021).

For each hybrid individual, we subsequently constructed histograms of relative coverages for LOH sites preserving either *C. ohridana* (O) or ‘ghost’-genome (X) alleles, with bin widths determined using the Freedman–Diaconis rule (Freedman and Diaconis 1981). We analyzed histograms for aforementioned biologically relevant values using a nonlinear least square method in Gnuplot, fitting them by gamma distributions that assumed a single type of event or mix of events, such as conversions and deletions simultaneously. The best model for each data set was selected through F-test comparisons of nested models, whose fit allowed us to quantify contributions of different processes causing observed LOHs in each hybrid’s dataset (Janko et al. 2021).

#### Detection of interspecific introgression of mtDNA

To estimate the gene flow between *C. ohridana* and their hybrids in mitochondrial compartment, we applied the Isolation with Migration MCMC models using IMa3 software v.1.13 (Hey et al. 2018). Using available *cyt b* mitochondrial data, we divided the sequences into two sets among which we estimated splitting time and migration; those from *Cobitis* sp. ODS hybrids and those of *C. ohridana*. The latter were subsequently split into two datasets, one including all available *C. ohridana* sequences and the other including only those from sites where *C. ohridana* is sympatric with *Cobitis* sp. ODS. After several preliminary short runs to figure out appropriate priors, we initiated a long MCMC run with an 10,000,000 burn-in step, and after burn-in, sampling every 100^th^ step, to collect 100,000 genealogies in total to reach enough genealogies for L-mode run. The MCMC analysis utilized 80 heated chains, following a standard approach, to provide adequate mixing across the parameter space. To further assess the migration dynamics, we used a series of likelihood-ratio tests for nested models and Chi-square test to evaluate their significance. The models tested included: isolation with no migration, migration only from *C. ohridana* to the hybrids, migration only from the hybrids to *C. ohridana*, and equal migration rates between the populations after their split.

#### Detection of nuclear interspecific introgression using exome-capture data

To investigate whether sequence variability of exons of nuclear genes in *C. ohridana* samples bear traces of interspecific gene exchange with the other potential parental species of Albanian hybrids, we employed the Dsuite v0.5 software (Malinsky et al. 2021) utilizing the ‘Dtrios’ module to generate D statistics (ABBA-BABA), Z scores, and f4 ratios. In order to determine whether potential gene exchange might have been mediated by asexual hybrids, we performed three separate analyses with distinct configurations in the ‘SETS’ input file.

First, we grouped individuals by species to let the algorithm estimate the overall levels of introgression. We ran this analysis twice, first using only *C. ohridana* specimens from localities where they are sympatric with *Cobitis* sp. ODS, and then with the individuals of *C. ohridana* from localities where they are allopatric to *Cobitis* sp. ODS.

Second, we treated each individual as a unique entity to capture individual-specific introgression patterns. To focus on introgression events between *C. ohridana* and other species, *C. ohridana* was consistently positioned as P3 in the phylogenetic setups, with different members from the *Cobitis* sensu stricto clade assigned to P1 and P2 positions (Fig. 4h). Each record in the dataset thus represents measurements obtained from specific combinations of individual organisms identified at positions P1, P2, and P3. To test whether *C. ohridana* samples which are sympatric and allopatric with *Cobitis* sp. ODS populations significantly differ in their estimated rate of introgression with *C.* sensu stricto, we used two pathways. First, we utilized Generalized Linear Mixed Models (GLMMs) to assess the impact of geographical (sympatric vs. allopatric) variations on D statistic, Z score, and f4 ratio values with individual identifiers at the P3 position treated as random effects to control for non-independence within the dataset. Additionally, we applied a block design to address non-independence of phylogenetic data further. This approach involved defining unique combinations of individuals at P1 and P2 positions as blocks. Within each block, we calculated median D statistics, Z scores, and f4 ratios for sympatric and allopatric groups of *C. ohridana*, and then performed Wilcoxon rank sum tests to compare these medians, thus assessing consistency of introgression across different genetic backgrounds of tested combination of individuals.

### Cytogenetic analyses

#### Mitotic chromosome preparation and karyotype analysis

Metaphase chromosomes were prepared according to Marta et al. (2020) with slight modifications. Briefly, fish were injected with 0.1% colchicine solution (1 ml/100 g of body weight) 45 min before being sacrificed using an overdose of 2-phenoxyethanol anesthetics agent (Sigma). The kidneys were removed, dissected in 0.075 M KCl, and treated hypotonically for 8 min in 0.075 M KCl. The resulting suspension was then fixed in fresh fixative (methanol: acetic acid 3:1, v/v), and spread onto slides. Metaphases of *C. ohridana* (n=12) and hybrid individuals (n=49) were inspected by conventional Giemsa or DAPI staining to confirm the number and morphology of their chromosomes, which were classified into four categories: metacentrics, submetacentrics, subtelocentrics, and acrocentrics according to Levan et al. (1964).

#### Meiotic pachytene chromosome preparation and immunofluorescent staining

Pachytene chromosomes were obtained for *C. ohridana* males and females as well as diploid and triploid *Cobitis* sp. ODS females according to protocols described in Dedukh et al. (2020). After manual homogenization, the suspension was placed in 1/3 of 1× PBS in the case of male gonads or 0.2 M Sucrose and 0.2% Triton X100 in the case of female gonads on SuperFrost® slides (Menzel Gläser). Afterward, chromosomes were fixed in 2% PFA, dried out, washed 1× PBS, and used for immunofluorescent staining.

Immunostaining was performed according to Dedukh et al. (2020) with the detection of lateral and central components of synaptonemal complexes by rabbit polyclonal antibodies against SYCP3 protein (dilution 1:200, ab14206, Abcam) and by chicken polyclonal antibodies against SYCP1 protein (dilution 1:300, a gift from Sean M. Burgess). Crossing over loci were detected by mouse monoclonal antibodies against mismatch repair protein 1 (MLH1) (dilution 1:50, ab15093, Abcam). Applied secondary antibodies included Alexa-594-conjugated goat anti-rabbit IgG (H + L) (concentration 1:200, A-11012, Thermo Fisher Scientific), Alexa Fluor™ 594 conjugated goat anti-chicken IgY (H+L) (concentration 1:200, A-11042, Thermo Fisher Scientific) and Alexa-488-conjugated goat anti-mouse IgG (H + L) (concentration 1:200, A-11001, Thermo Fisher Scientific). Primary and secondary antibodies were washed three times in 1× PBS with 0.05% Tween-20. After washing from secondary antibodies, slides were dehydrated in ethanol row (50%, 70% and 96%), dried, and mounted in Vectashield/DAPI (1.5 mg/ml) (Vector, Burlingame, Calif., USA).

#### Meiotic diplotene chromosome preparation

Diplotene chromosomal spreads (i.e. lampbrush chromosomes) were successfully prepared from adult *C. ohridana* and diploid *Cobitis* sp. ODS females following Dedukh et al. (2020). Vitellogenetic oocytes (0.8–1.5 mm in diameter) were taken from females within the OR2 saline (82.5 mM NaCl, 2.5 mM KCl, 1 mM MgCl_2_, 1 mM CaCl_2_,1mM Na_2_HPO_4_, 5 mM HEPES (4-(2-hydroxyethyl)-1-piperazineethanesulfonic acid); pH 7.4). Individual oocytes were transferred to ‘5:1’ medium (83 mM KCl, 17 mM NaCl, 6.5 mM Na_2_HPO_4_, 3.5 mM KH_2_PO_4_, 1mM MgCl_2_, 1 mM DTT (dithiothreitol); pH 7.0–7.2) to isolate nuclei. Nuclear envelopes were manually removed in one-fourth strength ‘5:1’ medium with the addition of 0.1% paraformaldehyde and 0.01% 1M MgCl_2_ in a chamber attached to a slide. Slides with oocyte nuclei contents underwent centrifugation for 20 min at +4°C, 4000 rpm, fixed for 30 min in 2% PFA in 1× PBS, dehydrated in ethanol row (50%, 70%, 96%), and dried.

#### CGH and FISH

A subset of 23 hybrid individuals with chromosomal spreads was selected for further comparative genome hybridization (CGH) analyses to identify parental chromosomal sets. Whole genomic DNA (gDNA) of one parental species *C. ohridana* and *C. elongatoides* – a representative of *Cobitis* sensu stricto clade as a parental species – was used for probe preparation. Individual gDNA samples were labeled with biotin-16-dUTP (Roche, Mannheim, Germany) and digoxigenin-11-dUTP (Roche) using a Nick Translation Mix (Roche) following the protocol supplied by the manufacturer.

For fluorescent *in situ* hybridization (FISH) on mitotic and meiotic chromosomes of *C. ohridana* and hybrids, we selected pancentromeric (SatCE04) and chromosome-specific (SatCE01) probes according to previously published data (Marta et al. 2020). gDNA of *C. elongatoides* was used for DNA probe amplification and labeling was performed by biotin-16-dUTP (Roche, Mannheim, Germany) and digoxigenin-11-dUTP (Roche) using PCR.

Before FISH and CGH, slides with mitotic chromosomes were additionally treated with 0.01% pepsin/0.01M HCl. The hybridization and detection procedure for CGH was carried out under conditions described in Majtánová et al. (2016). In the case of FISH, denaturation of probes was performed directly on slides in a hybridization mix (50% formamide, 10% dextran sulfate, 2× ЅЅС, 5 ng/μl labeled probe, and 10–50 fold excess of tRNA) at 77°C for 5 min following Marta et al. (2020). The biotin-dUTP labeled probes were detected using streptavidin Alexa 488 (Invitrogen, San Diego, Calif., USA). The digoxigenin-dUTP labelled probes were detected using Anti-Digoxigenin-Rhodamin (Roche). The chromosomes were counterstained with Vectashield/DAPI (1.5 mg/ml) (Vector, Burlingame, Calif., USA).

#### Cytogenetic image processing

Chromosomal preparations were examined by an Olympus Provis AX 70 epifluorescence microscope. Images of metaphase chromosomes were recorded with a cooled Olympus DP30BW CCD camera. The IKAROS and ISIS imaging programs (Metasystems, Altlussheim, Germany) were used to analyse grey-scale images. The captured digital images from GISH experiments were pseudocoloured (blue for DAPI, red for Anti-Digoxigenin-Rhodamin, green for Invitrogen FITC-Streptavidin) and superimposed using Microimage and Adobe Photoshop software, v. CS5, respectively.

### Visualization

*Cobitis* distribution map was prepared in Procreate v. 5.3.14. All figures were finalized in Adobe Photoshop v. CS5.

## Acknowledgements

We are grateful to Dovilė Barcytė and Pilar Ochoa for technical support, Danilo Mrdak, Nikola Hristovski, Iain Wilson for support in the field, and Jasmina Šandová for help with illustration. JV received support from the ES-TAF-2061 SYNTHESYS Project, financed by European Community Research Infrastructure Action under the FP7 “Capacities” Programme at the Museo Museo Nacional de Ciencias Naturales (CSIC). Computational resources were provided by the e-INFRA CZ project (ID:90254), supported by the Ministry of Education, Youth and Sports of the Czech Republic.

