## Supplementary Figures S1-S3 for "Zombie Gene Flow: Asexual Hybrids Mediate Extensive Genomic Introgression from Extinct Species Into Their Sexual Parent"

### Zombie Gene Flow: Asexual Hybrids Mediate Extensive Genomic Introgression from Extinct Species Into Their Sexual Parent

**Jasna Vukić, Anatolie Marta, Filip Matura, Daniel Kulik, Jan Eisner, Radek Šanda, Tomáš Tichopád, Lukáš Choleva, Anabel Perdices, Jan Roslein, Jan Kočí, Spase Shumka, Denik Ulqini, Jan Kotusz, Dmitrij Dedukh, Karel Janko**

**Figure S1.**

Summary of all known and putative reproductive pathways in *Cobitis* sp. ODS hybrid asexuals. This figure complements the information shown in Fig. 5 of the main text.

a-g) Meiotic spreads during pachytene stages;
h-i) Meiotic spreads during diplotene stages.
a, b, h) *C. ohridana* females;
c, d, i) diploid *Cobitis* sp. ODS hybrids;
e-g) triploid *Cobitis* sp. ODS hybrids.
Thick arrows indicate bivalents, thin arrows indicate univalents, arrowheads indicate crossing overs.
Note the formation of full bivalents sets in *C. ohridana* detected by the presence of crossing over sites (red staining in a) and the accumulation of central (SYCP1 protein, red staining in b) and lateral (SYCP3 protein, green staining in b) components of synaptonemal complexes.
Note the aberrant pairing between orthologous chromosomes in nonduplicated diploid hybrid's pachytene oocyte (c) and the detection of the centromeric region on chromosomes corresponding to 'ghost' genome in nonduplicated pachytene oocytes by FISH with satCE04 repeat.
Note two populations of pachytene oocytes in triploid hybrids, with 25 bivalents (e) visualized by the presence of crossing overs and with 25 bivalents and 25 univalents (f) detected by the accumulation of SYCP3 and SYCP1 proteins. FISH mapping of satCE04 repeat on pachytene oocytes with 25 bivalents and 25 univalents showed signal accumulation only on univalents corresponding to 'ghost' genome. Diplotene oocytes of *C. ohridana* (h) contain 25 bivalents with arrowheads indicating bivalent which is enlarged in (j). This bivalent was detected by chromosome specific probe (SatCE1, indicated by thick arrows) and shows weak signal from centromeric (SatCE4, indicated by thin arrows) probe.
Diplotene oocytes of diploid hybrid (i) contain 50 bivalents consisting of 25 *C. ohridana* bivalents and 25 bivalents corresponding to 'ghost' genome. (k, l) Enlarged bivalents from full diplotene chromosomal set from (i) visualized by chromosome specific probe (SatCE1, indicated by thick arrows) and centromeric (SatCE4, indicated by thin arrows) repeats. Bivalent corresponding to *C. ohridana* chromosome (k) showed weak accumulation of centromeric repeat (SatCE4) while bivalent formed by chromosomes corresponding to 'ghost' genome (l) exhibited much more intensive accumulation of centromeric repeat (SatCE4, indicated by thin arrows).

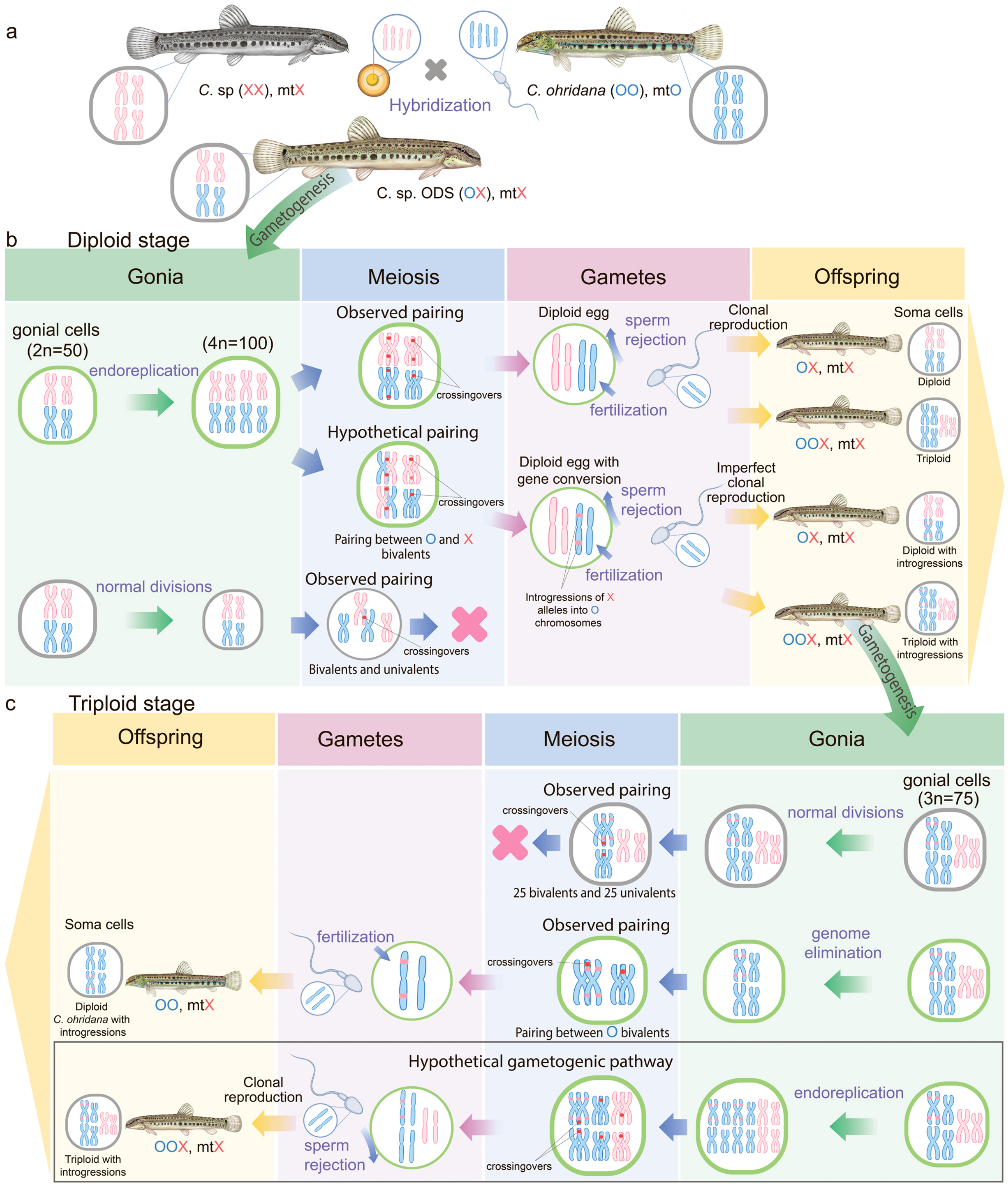
a
C. sp (XX), mtX
Hybridization
C. ohridana (OO), mtO
C. sp. ODS (OX), mtX
Gametogenesis
b
Diploid stage
Gonia
Meiosis
Gametes
Offspring
gonial cells (2n=50)
(4n=100)
endoreplication
Observed pairing
crossingovers
Hypothetical pairing
Pairing between O and X bivalents
Diploid egg
sperm rejection
fertilization
Clonal reproduction
Soma cells
OX, mtX
Diploid
OOX, mtX
Triploid
Diploid egg with gene conversion
Introgressions of X alleles into O chromosomes
Imperfect clonal reproduction
Diploid with introgressions
Triploid with introgressions
normal divisions
Bivalents and univalents
c
Triploid stage
gonial cells (3n=75)
25 bivalents and 25 univalents
genome elimination
Pairing between O bivalents
OO, mtX
Diploid C. ohridana with introgressions
Hypothetical gametogenic pathway

**Figure S2**.

Divergence time estimation and phylogenetic relationships among single representatives of each species in cytochrome *b* (a) and *S7* (b) genes. Trees are made ultrametric through application of uncorrelated clock in BEAST and translated to absolute time through calibration of Gibraltar strait opening (node highlighted by green star). Confidence bars for each node are indicated with horizontal bars. For non-supported nodes, bars are grey and ages in brackets.

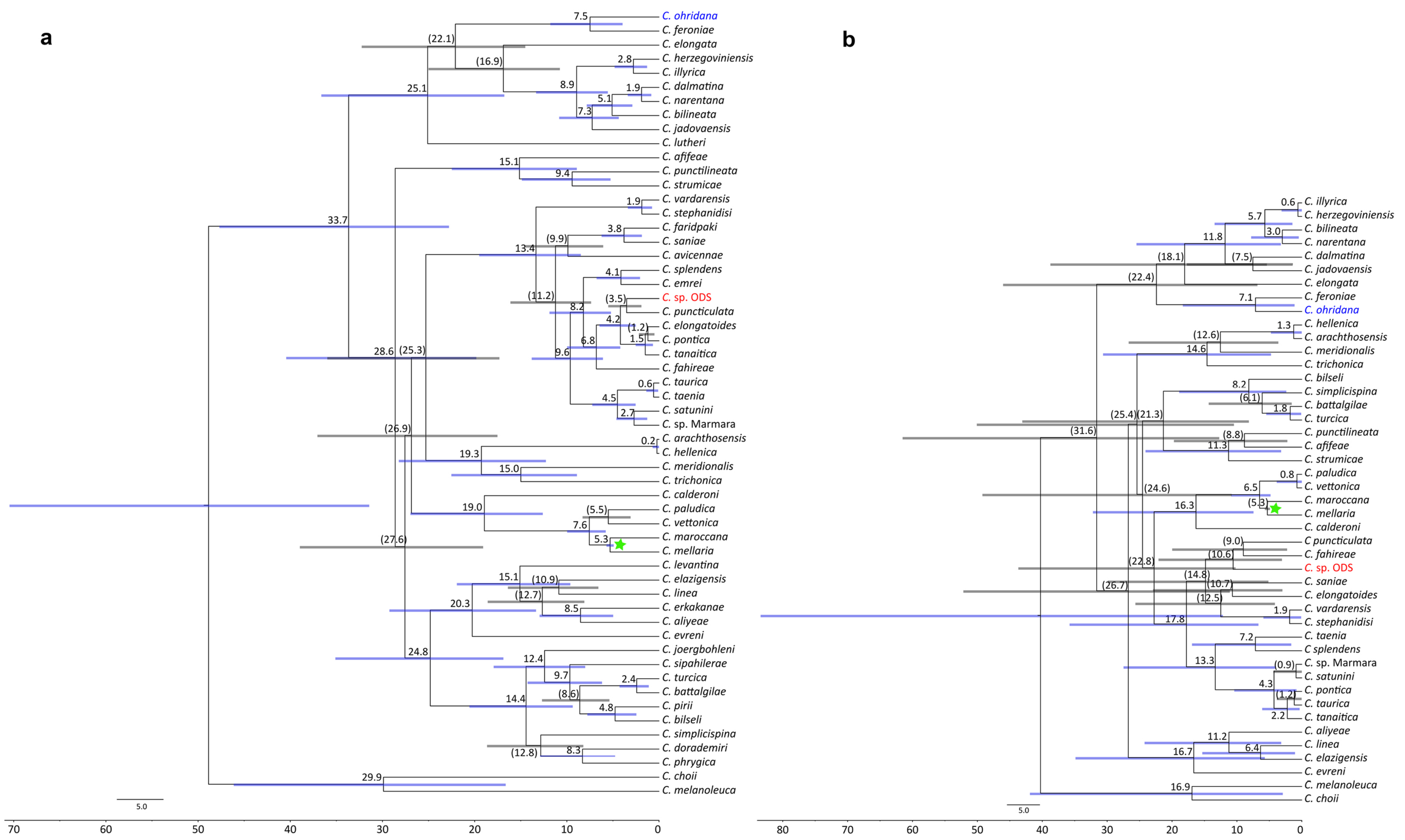
a
C. ohridana
C. feroniae
C. elongata
C. herzegoviniensis
C. illyrica
C. dalmatina
C. narentana
C. bilineata
C. jadovaensis
C. lutheri
C. afifeae
C. punctilineata
C. strumicae
C. vardarensis
C. stephanidisi
C. faridpaki
C. saniae
C. avicennae
C. splendens
C. emrei
C. sp. ODS
C. puncticulata
C. elongatoides
C. pontica
C. tanaitica
C. fahireae
C. taurica
C. taenia
C. satunini
C. sp. Marmara
C. arachthosensis
C. hellenica
C. meridionalis
C. trichonica
C. calderoni
C. paludica
C. vettonica
C. maroccana
C. mellaria
C. levantina
C. elazigensis
C. linea
C. erkakanae
C. aliyeae
C. evreni
C. joergbohleni
C. sipahilerae
C. turcica
C. battalgilae
C. pirii
C. bilseli
C. simplicispina
C. doradеmiri
C. phrygica
C. choii
C. melanoleuca
5.0
70
60
50
40
30
20
10
0
b
C. illyrica
C. herzegoviniensis
C. bilineata
C. narentana
C. dalmatina
C. jadovaensis
C. elongata
C. feroniae
C. ohridana
C. hellenica
C. arachthosensis
C. meridionalis
C. trichonica
C. bilseli
C. simplicispina
C. battalgilae
C. turcica
C. punctilineata
C. afifeae
C. strumicae
C. paludica
C. vettonica
C. maroccana
C. mellaria
C. calderoni
C puncticulata
C. fahireae
C. sp. ODS
C. saniae
C. elongatoides
C. vardarensis
C. stephanidisi
C. taenia
C splendens
C. sp. Marmara
C. satunini
C. pontica
C. taurica
C. tanaitica
C. aliyeae
C. linea
C. elazigensis
C. evreni
C. melanoleuca
C. choii
5.0
80
70
60
50
40
30
20
10
0

**Figure S3.**

Karyotypes and karyograms of *C. ohridana* (a, b) and diploid *Cobitis* sp. ODS (c-e). Both forms have 2n = 50 chromosomes. (a, b) *C. ohridana* has 16 m/sm chromosomes and 34 sb/a chromosomes. (c, d) *Cobitis* sp. ODS has one haploid set of *C. ohridana* chromosomes (8 m/sm and 17 st/a), and the other consisting of 22 m/sm and 3 st/a chromosomes belonging to ‘ghost’ parental species. m-metacentric, sm-submetacentric, st-subtelocentric, a-acrocentric chromosomes. (e) Metaphase plate with mapped SatCE4 allowing discrimination of ‘ghost’ genome. Scale bar = 10 µm.

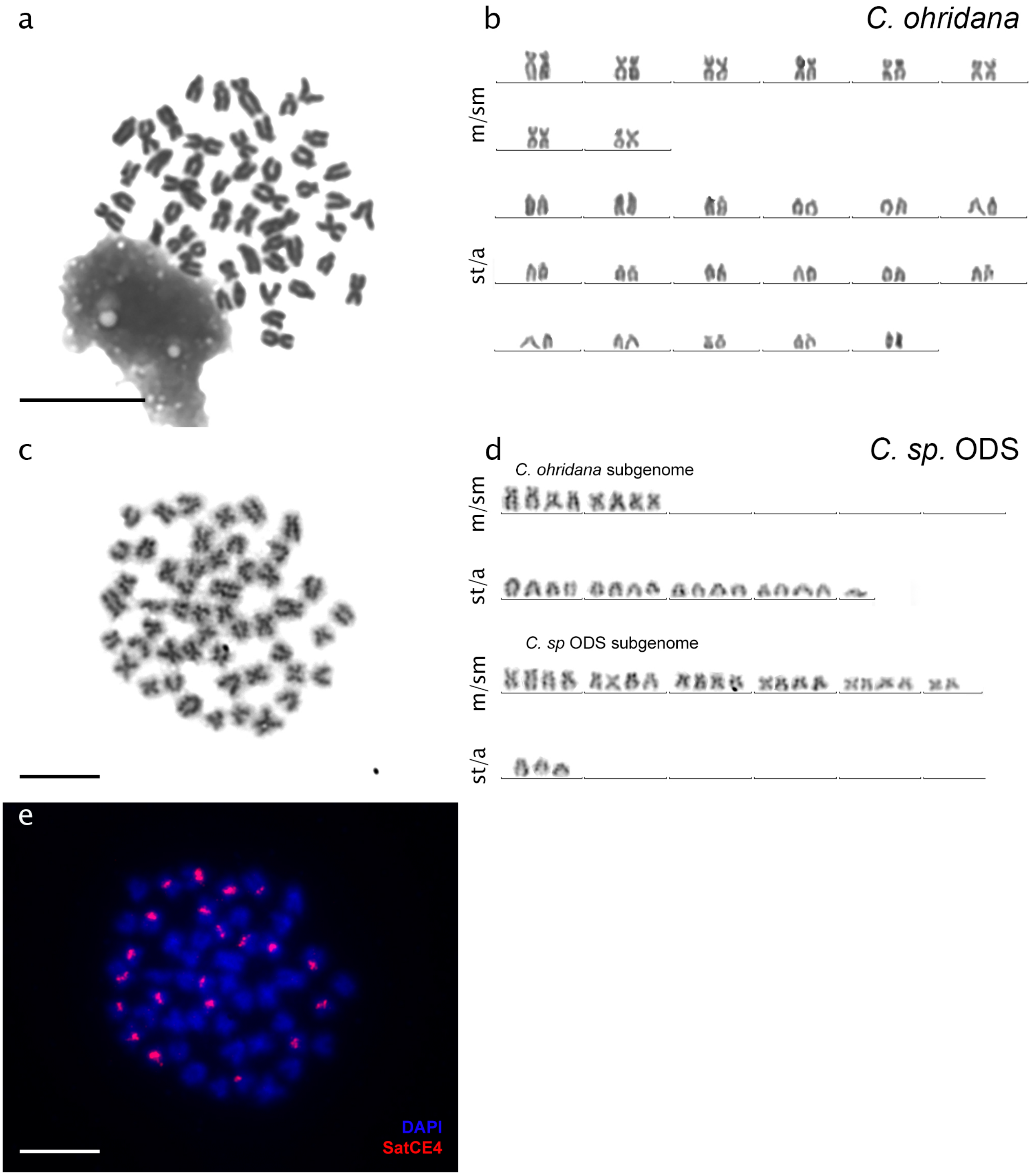
a
b
C. ohridana
m/sm
st/a
c
d
C. sp. ODS
C. ohridana subgenome
m/sm
st/a
C. sp ODS subgenome
m/sm
st/a
e
DAPI
SatCE4
